# In search of the Goldilocks zone for hybrid speciation

**DOI:** 10.1101/266254

**Authors:** Alexandre Blanckaert, Claudia Bank

## Abstract

Hybridization has recently gained considerable interest both as a unique window for observing speciation mechanisms and as a potential engine of speciation. The latter remains a controversial topic. It was recently hypothesized that the reciprocal sorting of genetic incompatibilities from parental species could result in hybrid speciation, when the hybrid population maintains a mixed combination of the parental incompatibilities that prevents further gene exchange with both parental populations. However, the specifics of the purging/sorting process of multiple incompatibilities have not been examined theoretically.

We here investigate the allele-frequency dynamics of an isolated hybrid population that results from a single hybridization event. Using models of 2 or 4 loci, we investigate the fate of one or two genetic incompatibilities of the Dobzhansky-Muller type (DMIs). We study how various parameters affect both the sorting/purging of the DMIs and the probability of observing hybrid speciation by reciprocal sorting. We find that the probability of hybrid speciation is strongly dependent on the linkage architecture (i.e. the order and recombination distance between loci along chromosomes), the population size of the hybrid population, and the initial relative contribution of the parental populations to the hybrid population. We identify a Goldilocks zone for specific linkage architectures and intermediate recombination rates, in which hybrid speciation becomes highly probable. Whereas an equal contribution of parental populations to the hybrid populations maximizes the hybrid speciation probability in the Goldilocks zone, other linkage architectures yield unintuitive asymmetric maxima. We provide an explanation for this pattern, and discuss our results both with respect to the best conditions for observing hybrid speciation in nature and their implications regarding patterns of introgression in hybrid zones.

**Summary:** Hybridization is observed ubiquitously in nature. Its outcome can range from extinction to the creation of new species. With respect to the latter, the probability of homoploid hybrid speciation, i.e. the formation of a new species as a result of an hybridization event without changes in the ploidy of the organism, is a hotly debated topic. Here, we analyze a minimal model for homoploid hybrid speciation, in which reproductive isolation is achieved by means of (postzygotic) Dobzhansky-Muller incompatibilities. When these postzygotic genetic incompatibilities are resolved in the hybrid population, their reciprocal sorting can result in reproductive isolation from both parental populations, thus creating a hybrid species. We show that, in accordance with the current literature, hybrid speciation tends to be rare. However, specific arrangements of the genes responsible for reproductive isolation can make reciprocal sorting almost unavoidable and thus create barriers to the parental population in an almost deterministic matter. We discuss the implications of these results for hybrid speciation and patterns of introgression in nature.

## Introduction

The role of hybridization in adaptation and speciation is an ongoing contentious question (Barton and Bengtsson, 1986; Rieseberg, 1997; Arnold et al., 1999; Buerkle et al., 2000; Barton, 2001; Mallet, 2007; Abbott et al., 2013; Servedio et al., 2013; Nieto Feliner et al., 2017; Schumer et al., 2018). On the one hand, hybridization may serve as a source of genetic variation. Various examples of adaptive introgression have been reported (reviewed in Hedrick, 2013), and it has been argued that hybridization may provide the fuel for adaptive radiations Seehausen (2013). On the other hand, gene flow between diverging populations may slow down or even reverse speciation either by purging isolating barriers or by one population swamping the other (Seehausen et al., 2008; Turissini et al., 2017). Thus, hybridization may act both as an engine of speciation and a boost to genetic variation, and as a detrimental mechanism that reduces population fitness and promotes extinction. This duality makes hybridization an important subject of study not only from an evolutionary but also a conservation biology point of view.

Hybrid speciation describes a scenario in which hybridization is essential for the formation of a “daughter” species that is isolated from both its parental species. The term “hybrid speciation” covers different scenarios that can be distinguished by the mechanism responsible for the buildup of reproductive isolation. In the case of polyploidization, the newly formed species consists of a fusion the genome of the two parents. The parents can be of the same species (autopolyploidization, although the hybrid species tends to be outcompeted by the parental diploid (Mallet, 2007)), or different ones (allopolyploidization), resulting in a single-step speciation event. In contrast, homoploid speciation (or recombinational speciation) corresponds to the formation of a hybrid species without a change in the ploidy level. This mechanism requires the existence of genetic barriers between the parental populations and the newly formed hybrid population, while still allowing the formation of sufficiently fit F1 hybrids. Despite this apparent paradox, numerous empirical cases have recently been reported (Schwarz et al., 2005; Mavárez et al., 2006; Larsen et al., 2010; Hermansen et al., 2011; Kang et al., 2013; Yakimowski and Rieseberg, 2014; Lamichhaney et al., 2018). Whether all of these represent true cases of homoploid hybrid speciation has been subject to debate. This debate has been led mainly around the definition of hybrid speciation and the resulting implications for the reported cases of empirical evidence (Schumer et al., 2014; Nieto Feliner et al., 2017; Schumer et al., 2018). However, to our knowledge there exists little work that has evaluated the probability of hybrid speciation theoretically.

Buerkle et al. (2000) studied a specific case of hybrid speciation via two overlapping parental inversions. Their simulations suggested a rather narrow parameter range in which hybrid speciation is possible, and indicated that (among other restrictions) high fertility of F1 hybrids is necessary to produce a stable hybrid population, which, as a consequence, is only poorly isolated from its parental species. Moreover, Schumer et al. (2015) studied the conditions for reciprocal sorting of genetic incompatibilities. A single genetic incompatibility can only isolate the hybrid population from one of its parental origins. However, if multiple DMIs exist between two species, in a hybrid population they might be resolved reciprocally with respect to the parental allelic origin, which can result in a hybrid species that is isolated from both parental populations. Proposing this model, Schumer et al. (2015) demonstrated via simulations that pairs of genetic incompatibilities can trigger hybrid speciation when there is a cost to both the ancestral genotype and the incompatibility-bearing genotypes. Based on a similar model, Comeault (2018) recently investigated the impact of adaptive loci linked to genetic incompatibilities on the probability of homoploid hybrid speciation in a stepping-stone model.

Inspired by Schumer’s model, we here provide a detailed analysis of the probability and dynamics of (reciprocal) sorting of classical Dobzhansky-Muller incompatibilities (DMIs; Bateson, 1909; Dobzhansky, 1936; Muller, 1942). A DMI consists of two (individually neutral or beneficial) alleles at different loci that are negatively epistatic, i.e., their combination is deleterious. The Dobzhansky-Muller model is arguably the most widely accepted model to explain the buildup of intrinsic postzygotic isolation in allopatric populations, and many examples of DMIs have been identified empirically (Presgraves, 2003; Kao et al., 2010; Corbett-Detig et al., 2013; Scarpino et al., 2013; Colomé-Tatché and Johannes, 2016; Ono et al., 2017; Marsit et al., 2017; Guerrero et al., 2017). Under this premise, such DMIs are expected to be the most prevalent type of genetic incompatibility that can be involved in the reciprocal sorting and thus contribute to hybrid speciation. However, as compared with the fitness landscape in Schumer et al. (2015), direct selection on the derived alleles is assumed to be absent or weak as compared with epistasis, and thus classical DMIs are asymmetric (Orr, 1995). We explain the differences between the models in Supplementary Section S 1.

We identify several parameters that greatly influence the probability of hybrid speciation via DMIs. Specifically, we quantify how the population size, the initial contribution of parental alleles, and the linkage architecture affect the probability of hybrid speciation. As linkage architecture, we define the relative position of the different loci involved in the hybrid incompatibilities that contribute to the species barriers (see also Fig. 1). Consistent with (Schumer et al., 2015), we define hybrid speciation as the successful reciprocal sorting of incompatibilities, independent of the amount of isolation they confer. We discuss both weak and strong isolating barriers and consider recessive and codominant DMIs (Turelli and Orr, 2000; Bank et al., 2012), which differ considerably in their sorting patterns. Our results indicate that the linkage architecture of the DMIs plays an essential role, such that a specific architecture can make hybrid speciation almost unavoidable, whereas exchanging only two loci may make hybrid speciation impossible for otherwise unaltered parameter values. Thus, we identify a Goldilocks zone of hybrid speciation, in which an interplay of population size, linkage architecture and the relative contribution of parental genomes can make hybrid speciation more likely than previously assumed.

**Figure 1:**
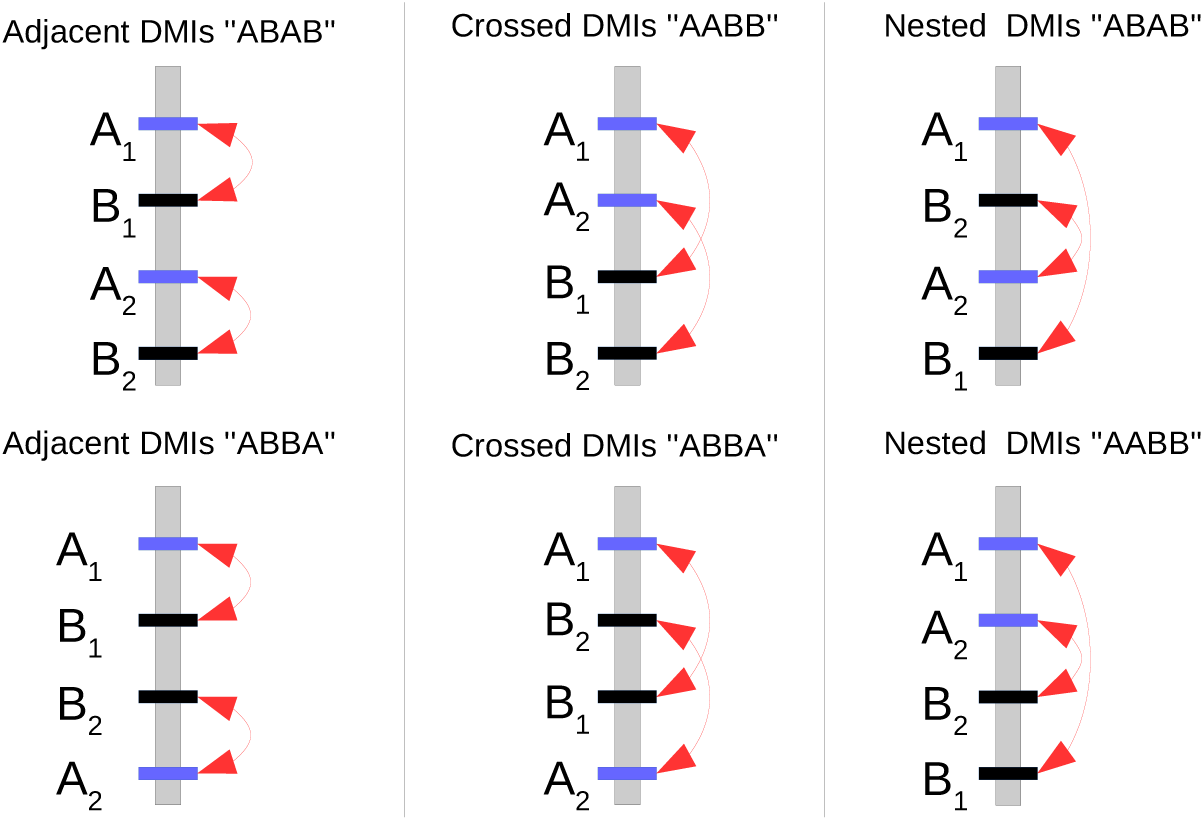
Illustration of all six different linkage architectures possible along a single chromosome. The *A_k_* loci are given in blue and *B_k_* in black. Red arrows show the incompatible interactions. The name of each architecture is derived from the architectures of the two incompatibilities and the order of the A and B loci.

## Results

### Model

We study the evolution of a single population of constant size *N* in discrete generations. We model four diallellic loci, *A*_1_, *A*_2_, *B*_1_, *B*_2_. At each locus, the derived allele is named after the locus, and the ancestral allele is named after its corresponding lower-case letter. Note that we do not detail here the two-locus model as it is fully included in the four-locus model. It can be obtained by keeping only loci *A*_1_ and *B*_1_. Derived alleles at the different loci are under direct selection (soft selection), with *α_k_* the (direct) fitness advantage of allele *A_k_* over *a_k_*, and *β_k_* the fitness advantage of allele *B_k_* over *b_k_*. Selection happens in the diploid phase of the life cycle. In addition, negative epistasis, *ϵ_k_*, which determines the strength of hybrid incompatibility, happens in a pairwise fashion between the derived *A_k_* and *B_k_* alleles (with *k* ∈ {1, 2}). Dominance affects only the epistatic interactions. In this manuscript, we focus mainly on two cases of dominance, which were proven representative of the general patterns (Bank et al., 2012): a recessive scenario and a codominant scenario, illustrated in Figure 2. We introduce 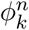 as a mathematical placeholder used to distinguish between the recessive and codominant scenario at the DMI *k*, with *n* the number of pairs of incompatible alleles in a genotype. Note that *n* = 1, *n* = 2 and *n* = 4 correspond to the *H*_0_, *H*_1_ and *H*_2_ incompatibilities in Turelli and Orr (2000). Therefore, for a codominant DMI, 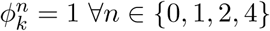 while for a recessive DMI, the effect of epistasis is masked for the double heterozygote genotype, i.e. 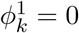 while ∀*n* ∈ {0, 2, 4}, 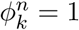.

**Figure 2:**
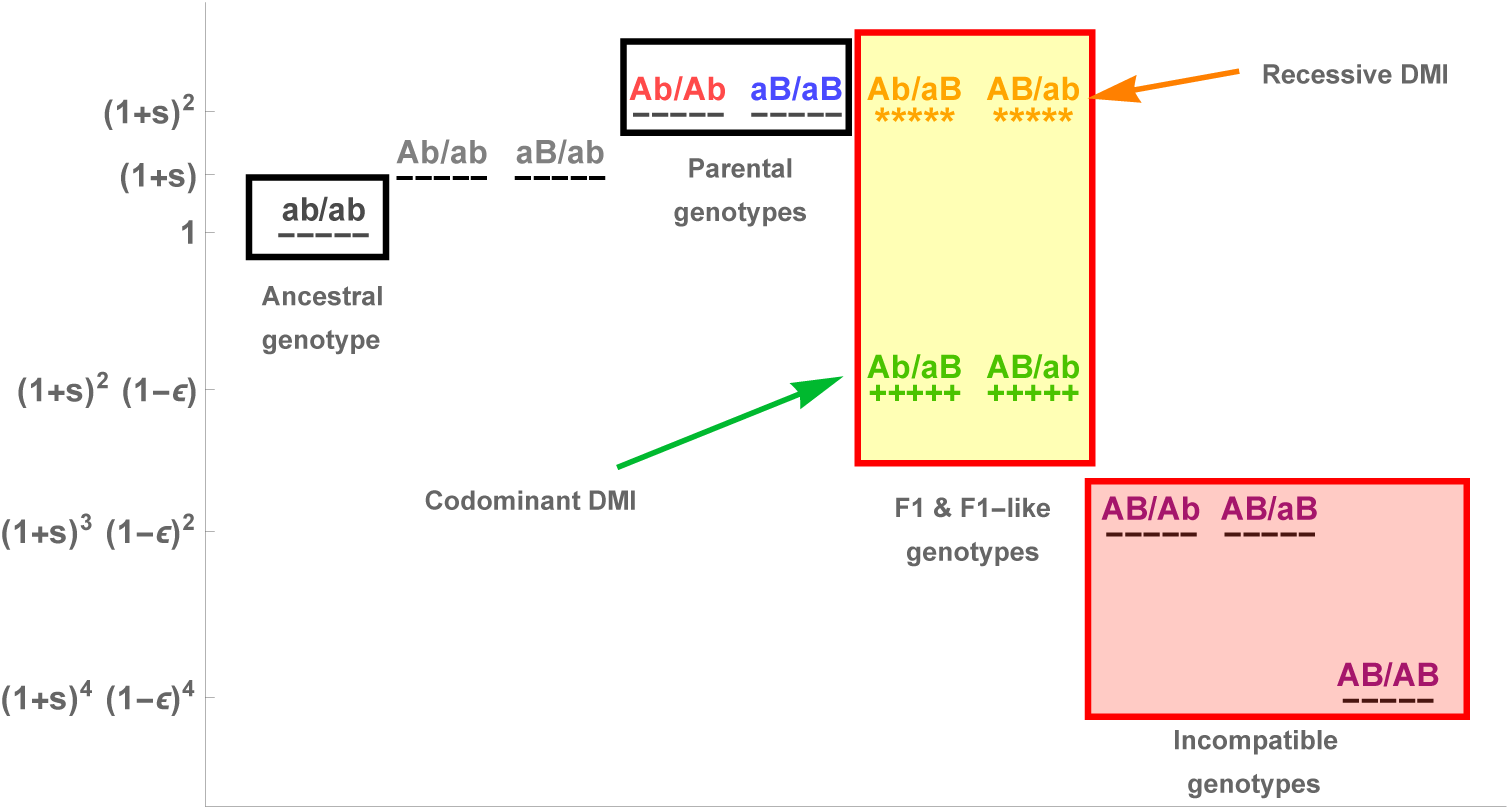
Fitness landscape of the 16 genotypes in the two-locus model, highlighting the effect of dominance of the incompatibility on the fitness of F1 hybrids. For simplicity, we illustrate the case of *α* = *β* = *s*. Note that there are only 10 genotypes represented here, as we do not distinguish the parental origins of each haplotype.

The population is initially composed of two single genotypes, since it is assumed to result from secondary contact between two monomorphic parental populations 1 and 2; *i_p_* denotes the contribution of the parental population 1 to the newly formed hybrid population. We assume that parental population 1 is fixed for the *A*_1_*b*_1_*A*_2_*b*_2_/*A*_1_*b*_1_*A*_2_*b*_2_ genotype and parental population 2 for *a*_1_*B*_1_*a*_2_*B*_2_/*a*_1_*B*_1_*a*_2_*B*_2_. The fitness of a genotype composed of haplotypes *i* and *j* is given by:

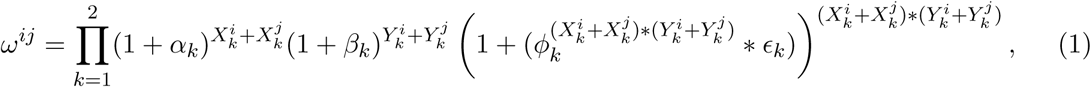
where 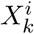 is the number of alleles *A_k_* in haplotype *i* and 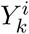 the number of alleles *B_k_* in haplotype *i*. The full fitness landscape is illustrated in Figure S1.

Mating is random. We assume that the parents generate an infinite pool of gametes from which zygotes are formed through multinomial sampling *M*(2*N*, *p*_1_, …, *p*_16_). Note that the deterministic case (i.e., in the absence of drift, *N* → ∞) can be obtained by skipping the multinomial sampling step during zygote formation.

As introduced above, hybrid speciation is defined as the fixation of a haplotype that is incompatible with both parental haplotypes, see Table 1. Indeed, if an individual homozygous for the *A*_1_*b*_1_*a*_2_*B*_2_ haplotype is backcrossed with an individual from, e.g., parental population 1, then the second DMI is expressed either in the F1 generation (codominant case) or in the F2 generation (recessive one). Similarly, introduction of an *A*_1_*b*_1_*a*_2_*B*_2_/*A*_1_*b*_1_*a*_2_*B*_2_ individual in the parental population 2 leads to the expression of the first DMI. This definition corresponds to an early stage speciation mechanism, leading to a hybrid population that is only partially isolated from both parental populations. Note that full isolation is impossible in this setting, as barriers responsible for full reproductive isolation would also prevent the formation of the hybrid population in the first place.

**Table 1:** Classification of possible haplotypes for the “Adjacent” linkage architecture.

|  |  |
| --- | --- |
| Ancestral haplotype | $a_1b_1a_2b_2$ |
| Parental pop. 1 haplotype | $A_1b_1A_2b_2$ |
| Parental pop. 2 haplotype | $a_1B_1a_2B_2$ |
| Hybrid haplotypes | $A_1b_1a_2B_2$ or $a_1B_1A_2b_2$ |
| “Epistasis-free” F2 haplotype | $A_1b_1a_2b_2$ or $a_1B_1a_2b_2$ or $a_1b_1A_2b_2$ or $a_1b_1a_2B_2$ |
| 1<sup>st</sup> incompatibility haplotypes | $A_1B_1a_2b_2$ or $A_1B_1A_2b_2$ or $A_1B_1a_2B_2$ |
| 2<sup>nd</sup> incompatibility haplotypes | $a_1b_1A_2B_2$ or $a_1B_1A_2B_2$ or $A_1b_1A_2B_2$ |
| Both incompatibilities haplotype | $A_1B_1A_2B_2$ |

We consider all possible linkage architectures formed by the two DMIs; they are illustrated in Figure 1. There are 6 different ways to arrange the 4 loci along a single chromosome (assuming the chromosome does not have an orientation). The two DMIs can be “Adjacent”, “Crossed”, or “Nested” (Fig. 1). Genetic distance between adjacent loci X and Y is given by 0 ≤ *r_XY_* ≤ 0.5. The distance between non-adjacent loci X and Y, separated by a single locus W, is given by *r_XY_* = *r_XW_* (1 − *r_WY_*) + *r_WY_* (1 − *r_XW_*). If the four loci are spread across multiple chromosomes, this represents a special case of the single chromosome scenario presented above, in which one or more recombination rates are set to 0.5. If not otherwise specified, we assume that all loci are located on different chromosomes, i.e. *r_XY_* = 0.5.

### Resolution of a single DMI

In the first part, we focus on the resolution of a single DMI following the formation of the hybrid population. With a single DMI, hybrid speciation according to our definition is impossible, because one of the negatively interacting partners in the DMI will invariably be lost, which makes the maintenance of a genetic barrier to both parental species impossible. Nevertheless, the study of a single incompatibility pair is necessary to understand which properties can be extrapolated to multiple incompatibilities. We characterize the resolution of the genetic conflict resulting from the contact between two diverged populations by quantifying the probability of fixation of the different haplotypes, the time of resolution of the DMI (i.e., the time until at least one of the incompatible alleles is lost) and the time to fixation of a single haplotype. In this section, we only focus on the loci *A*_1_ and *B*_1_ and drop the indices as they do not carry any information.

#### Dynamics following secondary contact

In a single randomly mating population such as the hybrid population we consider here, a DMI cannot be maintained unless directional selection is strong as compared with the epistatic effect of the incompatibility (Bank et al., 2012). This is because the formation of hybrid individuals initially leads to negative selection against both derived haplotypes. These haplotypes suffer from the incompatibility, either directly by forming an unfit hybrid genotype or indirectly through the production of unfit offspring. In contrast, the ancestral haplotype has an advantage as soon as it appears, and rises in frequency, because it only forms compatible genotypes and produces compatible offspring (if the proportion of incompatible *AB* haplotypes in the population remains low). As soon as the ancestral haplotype becomes frequent or either of the derived haplotypes becomes rare, this marginal advantage disappears, and the ancestral type will either be swamped by the more frequent derived type (in the case of direct selection acting on the derived alleles, i.e., if *α,β* > 0), or segregate neutrally (if *α, β* = 0). The incompatibility is usually resolved in favor of the more frequent derived allele (if they have similar fitness). Thus, one main determining factor is the initial frequency ratio between the two derived alleles (Supplementary Section S 2.1). Direct selection, as well as codominance of the incompatibility, reduce the impact of genetic drift (i.e., the outcome converges to the deterministic case). Indeed, once the DMI is resolved, selection increases the probability of fixation of a single derived allele (Haldane, 1927; Kimura, 1962). The codominance of the incompatibility shortens the time required to resolve the DMI (Fig. 3), and therefore reduces the time spent at low frequencies, where loss of the derived alleles because of drift is a likely outcome (see Supplementary Section S 2.2).

**Figure 3:**
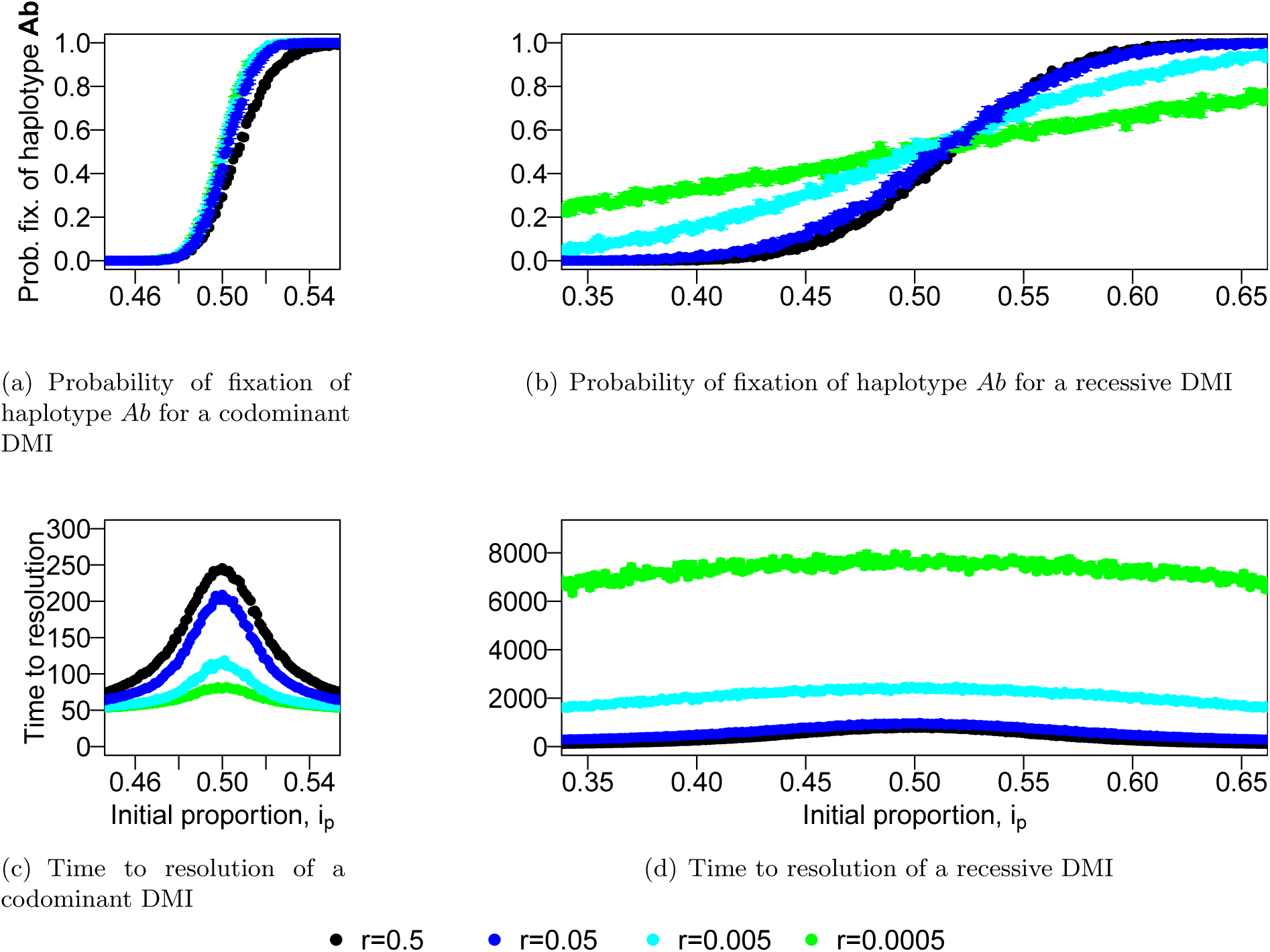
Recombination slows down the resolution of a codominant DMI while it speeds it up for a recessive one. We represent the probability of fixation of the *Ab* haplotype (top) for different recombination rates and different dominance schemes (codominant left, recessive right). We also illustrate (bottom) the time to resolve the genetic conflict (i.e., either allele *A* or *B* is lost). Each value is obtained from 1000 independent simulations. Note the much y-axis larger scale (x30) for panel d). Parameters used: *α* = *β* = 0.001, *ϵ* = −0.2, *N* = 5000.

#### Recombination has opposite effects under different dominance schemes

Recombination has a dual impact on the outcome of an hybridization event, depending on the dominance of the DMI, as illustrated in Figure 3 for haplotype *Ab*. Recombination breaks the association between the alleles of the parental haplotype and therefore leads to the formation of both the incompatible haplotype *AB* and the ancestral haplotype *ab*. On the one hand, this allows the expression of the incompatibility through the formation of the *AB* haplotype, which leads to faster sorting of the derived alleles. On the other hand, building a genotype with the ancestral haplotype protects both parental haplotypes from suffering from the genetic incompatibility, which leads to slower sorting of the derived alleles. The balance between these two effects is different between a recessive and a codominant DMI, which results in this opposite behavior, as illustrated in the following.

In the recessive case, recombination is necessary for the expression of the incompatibility. Thus, the need to form the incompatible haplotype overcomes any cost of generating the ancestral haplotype. An increase in recombination therefore always accelerates the resolution of a recessive DMI. This reduces the time the derived alleles spend at low frequency, which makes them less susceptible to being lost through genetic drift. This, in turn, reduces the probability that the ancestral haplotype becomes fixed.

In the codominant case, the incompatibility is already expressed in the F1 generation. Recombination is not necessary to express the incompatibility and therefore slows down the resolution of the DMI, as the ancestral haplotype prevents the effective purging of the parental haplotypes through the formation of *ab/Ab* or *ab/AB* individuals. In this situation, both derived alleles remain at a lower frequency much longer than in the recessive model, which makes them more likely to be both lost through genetic drift, resulting in the fixation of the ancestral haplotype.

### Resolution of two DMIs and hybrid speciation

Expanding from what we learned above, we now consider what happens when two incompatibilities exist between the parental populations. In contrast to the case of a single DMI, a new evolutionary outcome, namely hybrid speciation, becomes feasible with more than one DMI. As “hybrid speciation”, we denote the reciprocal sorting of the two DMIs, i.e. fixation of either alleles *A*_1_ and *B*_2_ or *A*_2_ and *B*_1_. Such a hybrid population will then be genetically isolated from both parental populations.

#### Isolation of the hybrid population by reciprocal sorting of two DMIs

Given the observed shape of the fixation probability of a derived allele in a single DMI case as a function of the initial contribution of both parental populations (Fig. 3), hybrid speciation should be observable only around a symmetric contact, and this condition should be more stringent for codominant incompatibilities than recessive ones (cf. Fig. 4). In Figure 4, we test this expectation by comparing the probability of hybrid speciation for two DMIs that are located on separate chromosomes (i.e., the “Adjacent” architecture from Fig. 1; colored dots in Fig. 4), with the expected probability of resolving two independent single DMIs for opposite derived alleles (e.g. first DMI resolved towards allele *A* and the second one for allele *B*; black dots). In the recessive case, the prediction for independent DMIs matches the hybrid speciation probability. In the codominant case, the independent expectation overestimates the probability of hybrid speciation. This can be explained by the faster resolution of the DMIs in the codominant model, which, even in the case of free recombination, leaves insufficient time for the two DMIs to become uncoupled and independently resolved, as the *A*_1_ and *A*_2_ loci start in maximum linkage disequilibrium. In the codominant case, this effect is amplified at low recombination rates as, in that case, the resolution of the DMIs happens even faster (Fig. S3), therefore preserving more of the initial linkage disequilibrium. This leads to a positive correlation between the fixation of the different *A_i_* alleles (as well as *B_j_* alleles), Figure S2. For the recessive case, the resolution of the two DMIs remains independent as it takes much longer to resolve the DMI.

**Figure 4:**
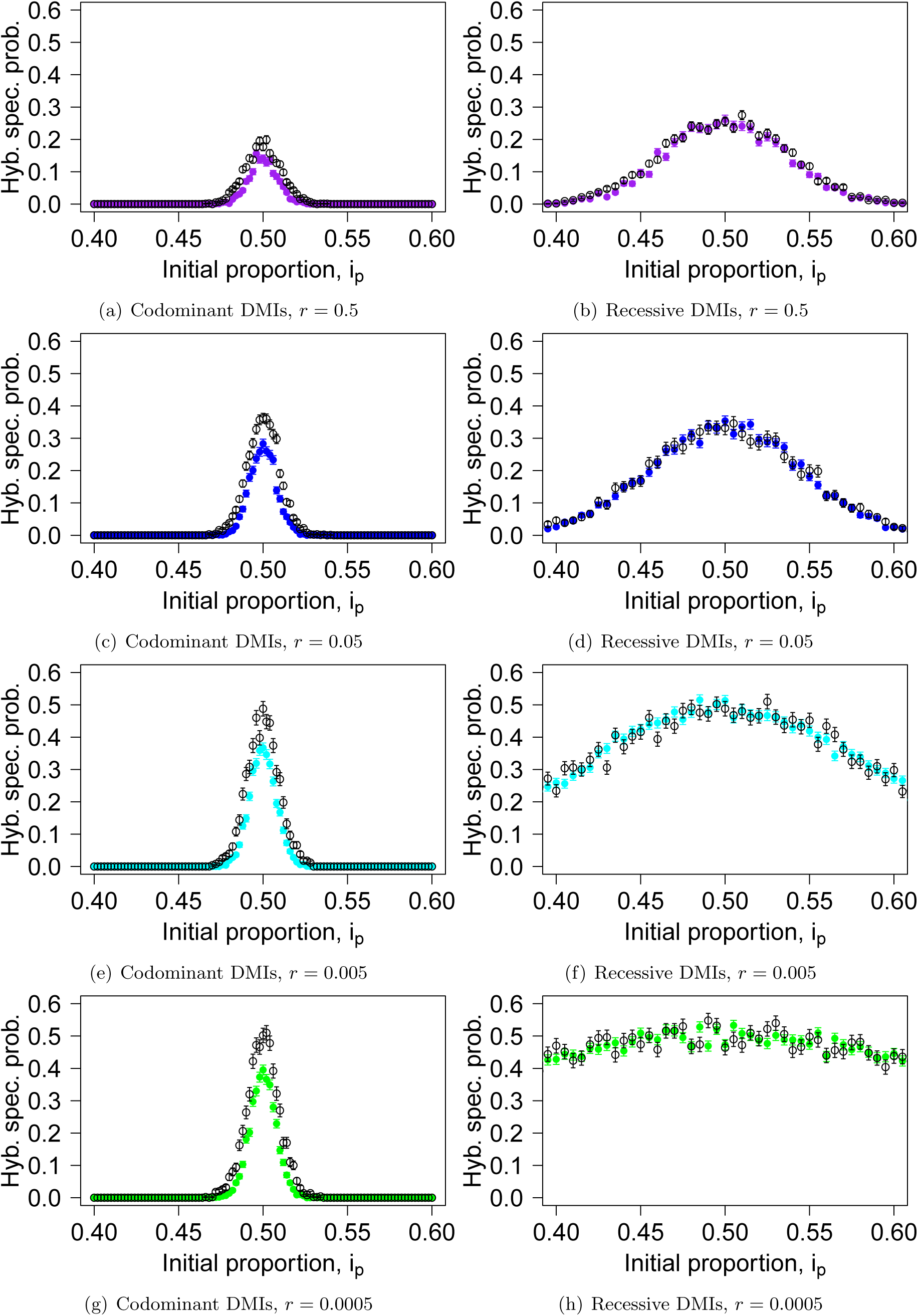
Hybrid speciation probability for codominant (left panels) and recessive (right panels) DMIs. The colored dots correspond to the probability of hybrid speciation for two DMIs situated on different chromosomes (*r*_23_ = 0.5). The genetic distance between the interacting loci is indicated below each panel (*r*_12_ = *r*_34_ = *r*). The black dots correspond to the predicted hybrid speciation probability based on the resolution of a single DMI. The fast resolution of the codominant DMIs leads to a correlation between their fate, which makes hybrid speciation less likely. Parameters used are *α_i_* = *β_i_* = .001, *N* = 5000, *ϵ* = −0.2. Each dot is obtained from 1000 replicates.

Figure S3 illustrates the mean time to resolve both DMIs in opposite directions conditioned on the outcome being hybrid speciation. Recombination has the same effect on the resolution of two DMIs than it did for a single one: it accelerates the resolution of recessive DMIs and slows down the resolution of codominant DMIs. However, the average resolution time is not affected by the initial proportion of the parental species; only trajectories that resolve slowly, which ensure that the initial linkage disequilibrium has been broken, can contribute to hybrid speciation, and we are conditioning on this outcome.

Due to the complexity of the dynamics, the possibility to obtain analytical results was very limited. However, we were able to obtain a condition for the (deterministic) local stability of the hybrid species with respect to back mutation or single migration events (see Supplementary Section S 4). Most notably, we find that the obtained stability condition is independent of the dominance of the incompatibilities.

#### The linkage architecture determines which alleles survive

Figure 5 illustrates the effect of recombination and the linkage architecture on hybrid speciation, when all loci are on the same chromosome (as opposed to one DMI per chromosome, as illustrated in Figure 4). As mentioned above, for codominant DMIs, recombination, on the one hand, allows the formation of the hybrid haplotype and helps to reduce the initial linkage disequilibrium. On the other hand, it slows down the resolution of the DMI through the formation of compatible haplotypes. Depending on the balance between these two effects, recombination impacts the probability of hybrid speciation differently.

**Figure 5:**
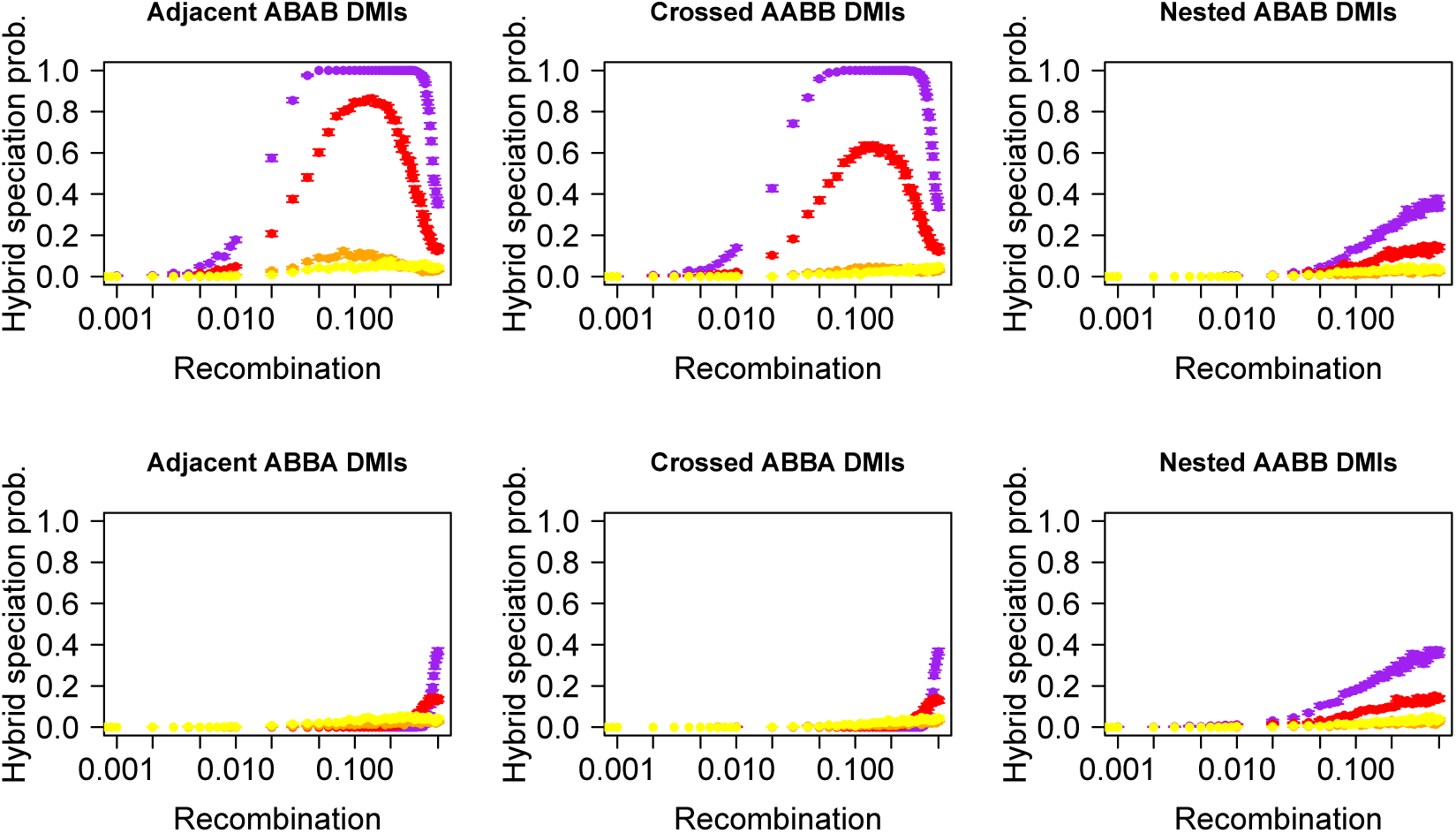
Hybrid speciation probability is a nonlinear function of recombination. We consider that all four loci have the same selective advantage (*α_k_* = *β_j_* = .001) and are equidistant along a single chromosome. The hybrid speciation probability is plotted for different population sizes: yellow corresponds to *N* = 50, orange to *N* = 500, red to *N* = 5000 and purple to *N* = 50000. Epistasis (*ϵ* = −0.2) is here codominant but we obtain qualitatively similar results for recessive incompatibilities, see Figure S9. The contribution of both parental populations here is symmetric (*i_p_* =0.5).

Assuming a symmetric contact, we observe that two of the linkage architectures, the “Adjacent ABAB” and “Crossed AABB” architectures, which have in common that the loci *A*_1_ and *B*_2_ are located at the ends of the chromosomal region that contains the four focal loci, exhibit a non-monotonic behavior with maximum hybrid speciation probability for intermediate recombination rates. Hybrid speciation becomes almost certain for large population size and indeed corresponds to the deterministic outcome (i.e. in the absence of drift) for these two linkage architectures. More precisely, we observe a local maximum of the hybrid speciation probability for recombination rates around *r* = 0.1. The “Adjacent ABAB” and “Crossed AABB” architectures, which show this behavior, are characterized by a higher marginal fitness of the *A*_2_ and *B*_1_ alleles compared to the other alleles in the deterministic case, which promotes hybrid speciation. For all other architectures either *A*_1_ and *A*_2_ or *A*_2_ and *B*_2_ have the highest marginal fitness. The higher fitness stems from the production of the *a*_1_*B*_1_*a*_2_*b*_2_ and *a*_1_*b*_1_*A*_2_*b*_2_ haplotypes (for the “Adjacent ABAB” architecture) that are relatively free of epistasis in the F2 generation. The outcomes of a single recombination event per genome for all 6 architectures are given in Table 2 and illustrates how the “Adjacent ABAB” and “Crossed AABB” architectures stand out in the production of the haplotypes that are needed of hybrid speciation. Importantly, recombination is necessary to generate these haplotypes, but too much recombination will cancel their advantage. Indeed, for *r* = 0.5, all haplotypes are produced at the same frequency in the absence of selection. This dual effect of recombination leads therefore to the observed maximum in the hybrid speciation probability for intermediate recombination rates. When the DMIs are located on two different chromosomes (as in Fig. 4), this effect does not appear. Indeed, while recombination still breaks linkage disequilibrium, it no longer generates the relatively “epistasis-free” haplotype and therefore leads to a monotonous increase in the hybrid speciation probability with increasing recombination rate. This behavior, specific to the “Adjacent ABAB” and “Crossed AABB” linkage architectures is observed both for codominant and recessive DMIs.

**Table 2:** Haplotypes produced in the F2 breakdown, assuming a single recombination event, explain how different linkage architectures leads to different outcomes for the same loci. By identifying the relatively “epistasis-free” haplotype formed, one can infer whether hybrid speciation may be a likely outcome. In blue, we highlight these “epistasis-free” haplotypes that are important for hybrid speciation and in yellow those that are important for fixation of the parental haplotype from population 1.

| Genetic architecture | F1 hybrid | Single recombination event between: |  |  |
| --- | --- | --- | --- | --- |
|  |  | loci 1 and 2 | loci 2 and 3 | loci 3 and 4 |
| Adjacent ABAB | | $A_1 B_1 a_2 B_2$<br>$a_1 b_1 A_2 b_2$ | $A_1 b_1 a_2 B_2$<br>$a_1 B_1 A_2 b_2$ | $A_1 b_1 A_2 B_2$<br>$a_1 B_1 a_2 b_2$ |
| Adjacent ABBA | | $A_1 B_1 B_2 a_2$<br>$a_1 b_1 b_2 A_2$ | $A_1 B_1 b_2 a_2$<br>$a_1 b_1 B_2 A_2$ | $A_1 b_1 b_2 a_2$<br>$a_1 B_1 B_2 A_2$ |
| Crossed AABB | | $A_1 a_2 B_1 B_2$<br>$a_1 A_2 b_1 b_2$ | $A_1 A_2 B_1 B_2$<br>$a_1 a_2 b_1 b_2$ | $A_1 A_2 b_1 B_2$<br>$a_1 a_2 B_1 b_2$ |
| Crossed ABBA | | $A_1 B_2 B_1 a_2$<br>$a_1 b_2 b_1 A_2$ | $A_1 B_2 b_1 a_2$<br>$a_1 b_2 B_1 A_2$ | $A_1 b_2 b_1 a_2$<br>$a_1 B_2 B_1 A_2$ |
| Nested AABB | | $A_1 a_2 B_2 B_1$<br>$a_1 A_2 b_2 b_1$ | $A_1 A_2 B_2 B_1$<br>$a_1 a_2 b_2 b_1$ | $A_1 A_2 b_2 B_1$<br>$a_1 a_2 B_2 b_1$ |
| Nested ABAB | | $A_1 B_2 a_2 B_1$<br>$a_1 b_2 A_2 b_1$ | $A_1 b_2 a_2 B_1$<br>$a_1 B_2 A_2 b_1$ | $A_1 b_2 A_2 B_1$<br>$a_1 B_2 a_2 b_1$ |

As illustrated in Figure S9, the recessive case is qualitatively similar to the codominant one. We recover the distinctive pattern between linkage architectures, where the “Adjacent ABAB” and “Crossed AABB” architectures are more likely to generate hybrid speciation for intermediate recombination rates. However, for the “Adjacent ABBA” and “Crossed ABBA” linkage architectures, the recessive case differs from the codominant by the existence of two local maxima for the hybrid speciation probability as a function of recombination. These two architectures are characterized by an indirect selective advantage of one of the two parental haplotypes over the other, as the partially derived haplotypes *A*_1_*b*_1_*b*_2_*a*_2_ and *a*_1_*b*_1_*b*_2_*A*_2_ are more likely to form than their counterparts (*a*_1_*B*_1_*b*_2_*a*_2_ or *a*_1_*b*_1_*B*_2_*a*_2_, see Table 2), which leads to a slightly higher marginal fitness of the *A*_1_*b*_1_*b*_2_*A*_2_ haplotype compared to *a*_1_*B*_1_*B*_2_*a*_2_. The first maximum is obtained at large intermediate recombination rates; it corresponds to the one observed for codominant DMIs. However, a second one can be observed at lower recombination rate if the population size reaches certain sizes. It results from a subtle balance between drift, recombination and selection, which we explain in detail in the Supplementary Section S 3.1.

The hybrid speciation probability for codominant versus recessive DMIs differs significantly when considering lethal incompatibilities, (see Supplementary Section S 3.2). Hybrid speciation becomes impossible for codominant DMIs because no viable hybrids can be produced. This is not the case for recessive incompatibilities, as they can partially escape the strong selection against hybrids. In fact, due to the masking effect provided in F1 and F1-like genotypes, we observe an almost indistinguishable pattern for deleterious (*ϵ* = −0.2) and lethal (*ϵ* = −0.99) recessive DMIs, Figure S13. Similarly, the time to hybrid speciation is similar between the deleterious and lethal recessive cases, Supplementary Section S 3.3.

Figure 5 also illustrates the impact of the population size on the outcome. In general a larger population size results in a higher probability of hybrid speciation. This is especially true when the deterministic outcome corresponds to hybrid speciation (i.e. the “Adjacent ABAB” and “Crossed AABB” architectures). Derived alleles are less likely to be lost during the reciprocal sorting of the genetic incompatibilities. The main exception to this rule exists when the deterministic outcome is the fixation of one parental haplotype. In that case, an intermediate population size will maximize the likelihood of hybrid speciation, as illustrated in Figure 5 for the “Adjacent ABBA” and “Crossed ABBA” architectures (and Fig. S9). This intermediate value corresponds to a balance between a strong drift regime in which the ancestral and “epistasis free” haplotypes are most likely to fix, and the deterministic regime in which the *A*_1_*b*_1_*b*_2_*A*_2_ parental haplotype fixes.

#### Symmetric contact is not always the best condition for hybrid speciation

Figure 5 was obtained for *i_p_* =0.5, i.e. when both parental populations contribute equally to the hybrid population. It corresponds to the case that is the most frequently investigated in models. Figure 6 illustrates what happens when we release this assumption. From the single-DMI dynamics, one would expect a decrease in the hybrid speciation probability as illustrated in Figure 5. This is not always true. Depending on the linkage architecture, the probability of hybrid speciation may be higher for asymmetric contributions from the parental populations. This phenomenon is also observed for intermediate recombination rates; thus, only a consideration of dominance scheme, recombination rate, and symmetry together allows for an accurate statement on the hybrid speciation probability (see Fig. 6). Table 2 provides us with an explanation for the observed pattern: for intermediate recombination rate (*r* ≈ 1/3), there is on average one recombination event per haplotype. For the two architectures concerned (“Adjacent ABBA” and “Crossed ABBA”), in this scenario and with perfect symmetry, both alleles *A*_1_ and *A*_2_ have a marginal fitness that is slightly higher than alleles *B*_1_ and *B*_2_ (Fig. S4), which leads to the fixation of the parental haplotype *A*_1_*b*_1_*b*_2_*A*_2_ in the deterministic case. Therefore, a lower initial frequency of these alleles at the initial contact balances this selective advantage, which results in higher hybrid speciation probabilities than under symmetry. This behavior is only observed for the two architectures discussed above (“Adjacent ABBA” and “Crossed ABBA”). For all other architectures, the two derived alleles that have a slight indirect selective are *A*_2_ and *B*_1_ for “Adjacent ABAB” and “Crossed AABB” (which correspond to the cases of high probabilities of hybrid speciation) or *A*_2_ and *B*_2_ for the two “Nested” architectures. In both cases, since the symmetry between the A and B alleles is respected, hybrid speciation is most likely at *i_p_* = 0.5.

**Figure 6:**
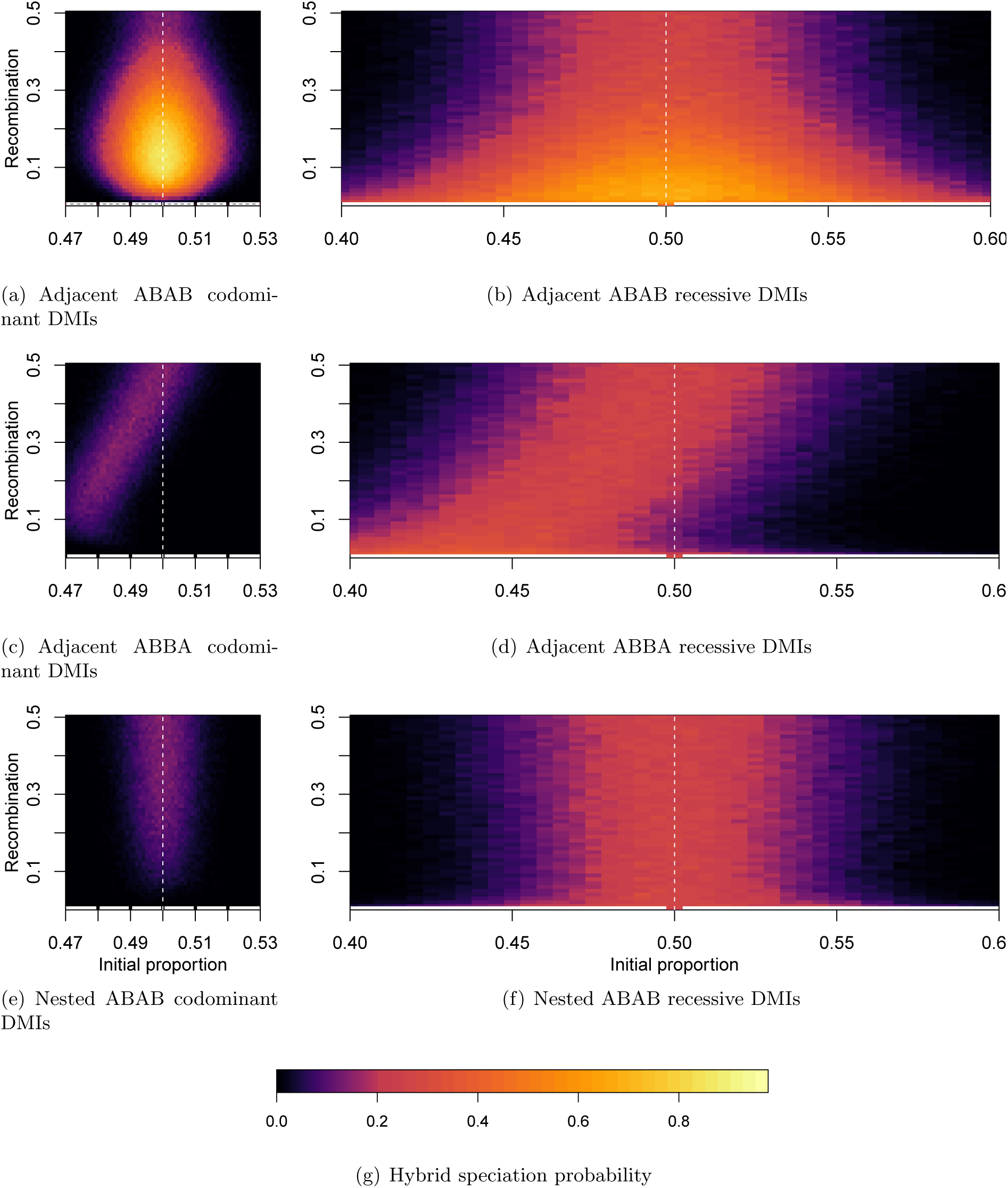
Probability of hybrid speciation for both recessive and codominant DMIs as a function of the genetic distance and the initial contribution of both parental species. Different linkage architectures generate an unexpected pattern: for the “Adjacent ABAB” architecture, we observe a Goldilocks zone for hybrid speciation; for the “Adjacent ABBA”, the probability of hybrid speciation is no longer symmetric along the *i_p_* = 0.5 axis (white dashed line).

## Discussion

We here characterized the purging process of single and multiple DMIs upon formation of a hybrid population. Specifically, we quantified the effects of the linkage architecture and the dominance of the epistatic interactions on the reciprocal sorting of incompatibilities, which has been proposed as a mechanism to induce homoploid hybrid speciation. We found that for the exact same set of loci, their order along the chromosome can increase the probability of observing hybrid speciation from unlikely to almost certain. We demonstrate that the main determinant of this pattern is which haplotypes are formed during the F2 breakdown. For the linkage architectures that promote hybrid speciation, there exists a Goldilocks zone in which an intermediate recombination rate maximizes the hybrid speciation probability. In addition, we show that symmetric contact of incompatible loci that are under equal selection pressure does not always generate the highest probability of hybrid speciation, and that this result cannot be predicted from the study of independent DMIs. Finally, for recessive DMIs, in which the F1 generation does not suffer a fitness disadvantage, reciprocal sorting is similarly probable with intermediate and strong epistasis. Conversely, hybrid speciation with lethal codominant DMIs is impossible.

### Linkage architecture as major determinant of hybrid speciation probability

Abbott et al. (2013) recently emphasized that “an important challenge in studies of hybrid speciation is to ask whether there is an ‘optimal’ genetic distance for homoploid hybrid speciation (Arnold et al., 1999; Gross, 2012).” Although Abbott et al. (2013) were arguably referring to the degree of divergence and thus, to the number of DMIs that have established between two species, our study adds an additional important factor to their list: the linkage architecture of the isolating barriers, and the recombination rate between them. Specifically, our results demonstrate that intermediate recombination rates and specific linkage architectures maximize the probability of hybrid speciation.

We can speculate whether the presence of multiple DMIs should increase the probability of hybrid speciation. Based on our results, we believe that it should depend on the nature of the incompatibilities: additional recessive DMIs should increase the hybrid speciation probability while more codominant DMIs should reduce it. Firstly, we showed that for codominant DMIs, the fixation of derived alleles of the same parental population is correlated (Fig. S2), which hinders their reciprocal sorting even if they are located on different chromosomes. Hence, recombination is not sufficient to decrease the initially existing linkage disequilibrium. Adding additional codominant incompatibility loci will result in stronger selection against F1 individuals which strengthens the correlation of parental alleles, and reduces the probability of reciprocal sorting. Secondly, we have shown that for lethal codominant DMIs (Fig. S13), hybrid speciation is impossible. Extrapolating from these two observations, we propose that this effect will outpace the increase in hybrid speciation probability due to having more chances to have at least one pair of reciprocal sorting.

On the other hand, in the recessive model, F1 hybrids do not suffer a fitness cost, and the fixation of derived alleles from the same parental origin is uncorrelated. In addition, stronger epistasis does not affect the probability of hybrid speciation in the recessive model (Fig. S9 and S13). Thus, having more than two recessive DMIs should increase the chances that at least two are reciprocally sorted, which is sufficient for hybrid speciation according to our definition.

Our results can be discussed in the context of parapatric speciation and the role of genomic islands of divergence (Via, 2012; Feder et al., 2012). According to the respective theory (Wu, 2001), during the speciation process and in presence of gene flow, islands of divergence are formed around the first genes involved in reproductive isolation. These genes will reduce gene flow locally around them, which favors the accumulation of weakly locally adapted mutations in their vicinity, as well as incompatible genes, reinforcing and extending islands of divergence. For hybrid speciation according to our model, the existence of such islands implies that many derived alleles *A_k_* may be found together on the same island, as locally adaptive loci tend to be rearranged in clusters (Yeaman, 2013). In such cases the linkage disequilibrium between the different *A_k_* alleles will be harder to break, which makes reciprocal sorting less probable. From the perspective of this model, hybrid speciation should therefore be most likely early during speciation, when no strong islands of divergence have formed yet. We further note that our model considers a single hybridization event without any further gene exchange with both parental populations, which resembles the colonization of a new environment. Continuous gene flow between populations, which is often a key feature of studies of genomic islands of speciation, should further reduce the probability of hybrid speciation (Comeault, 2018), because migration creates selective pressure against the hybrid haplotypes.

### The probability of hybrid speciation and reciprocal sorting in nature

Our results imply that while specific linkage architectures may indeed induce hybrid speciation with high probability, it remains on average unlikely, which is consistent with the scarcity of putative cases of homoploid hybrid speciation observed in nature (Schwarz et al., 2005; Mavárez et al., 2006; Larsen et al., 2010; Hermansen et al., 2011; Kang et al., 2013; Yakimowski and Rieseberg, 2014; Lamichhaney et al., 2018). Recently, Runemark et al. (2018) reported that the Italian sparrow hybrid species resulted from multiple occurrences of hybridization between the Spanish and House sparrow along the Mediterranean Sea, which is in concordance with our observation of a Goldilocks zone. The observed resolution of all hybridization events towards a single mitochondrial origin (i.e. all Italian sparrows possess the mitochondrial DNA of House sparrows) indicates that either the mitochondria play an important role with respect to the sorting of the incompatibilities, or that there is an asymmetry in the formation of the different hybrid populations, in which Spanish sparrow males mate with House sparrow females.

### The nature of genetic incompatibilities

Both theoretical considerations and empirical evidence suggest that most DMIs should be recessive (Orr, 1993; Presgraves, 2003). However, any kind of dominance pattern of the epistatic interactions can in theory exist (Coyne and Orr, 2004). Here, we showed that single codominant DMIs are resolved much faster than recessive ones. Therefore, when non-equilibrium populations are sampled, the excess of recessive incompatibilities may not necessarily reflect the true proportion of recessive incompatibilities but rather a sampling bias.

### The time to hybrid speciation

We specifically evaluated the timing of two events during the process of hybrid speciation. The first is when the conflicts caused by the DMIs are resolved by losing one of the interacting partners in each incompatibility pair; we call this the resolution time. From this point onwards, there are no epistatic interactions left in the population and the evolution of the remaining genotypes is dictated only by genetic drift and direct selection. The second event is the time at which all polymorphism is lost; conditioned on reciprocal sorting we call this the time to hybrid speciation. Only then, the isolating barrier to the parental populations is fixed in the population.

We find that initially, codominant DMIs are resolved faster, which leads to a short resolution time. This process is even faster with low recombination, because the lack of recombination decreases the probability to break the epistatically interacting haplotypes. (This also leads to a low probability of hybrid speciation for codominant DMIs with low recombination.) However, after resolution of the DMI, the derived alleles still need to become fixed in order for hybrid speciation to occur. This process is usually faster for recessive DMIs and compensates for the slower initial sorting, such that both codominant and recessive DMIs lead to similar times to hybrid speciation (Fig. S14 and S15).

Schumer et al. (2015) emphasized that hybrid speciation can sometimes happen quickly. Somewhat contradictory, we find that the time to hybrid speciation increases with smaller population sizes, and that there is relatively little variation in the times to hybrid speciation (Fig. S14 and S15). For example, for a population of size N=50, the time to hybrid speciation is around 5N=250 generations (i.e., longer than the average fixation time of a neutral allele) independent of the linkage architecture of the DMI. For a population of size N=50000, it is only around 0.1N=5000 generations. That is because both epistatic and direct selection act more efficiently in large populations. Therefore, the short time to reproductive isolation reported in the empirical examples (e.g., Lamichhaney et al., 2018) may reflect the size of the hybrid population rather than an inherent property of hybrid speciation.

### Population size and selection

In our model, we consider populations of constant size. Relaxing this assumption (i.e. switching from soft selection to hard selection), one would expect hybrid speciation to be less likely for at least two reasons. First, selection against the different derived alleles in the early purging phase is stronger; indeed with soft selection the effect of a mutation is weighted by the mean fitness of the population. Therefore, in maladapted populations, the effect of deleterious mutations is slightly dampened. Second, the expected decrease in population size that is associated with the purging phase increases the impact of drift, which means that reciprocal sorting is less likely even with favorable linkage architectures (as selection is not strong enough to counteract its effect). Lastly, even if the DMIs are resolved in opposite directions, the different derived alleles will be at low frequency when their interacting partner is lost, and therefore far more likely to be lost by drift subsequently. Overall, this implies that hybrid speciation via reciprocal sorting is on average less likely than illustrated here. Furthermore, this kind of contact between two diverged populations (or species) is usually geographically restricted, and therefore happens for small populations. However, this apparent rarity of hybrid speciation can be counteracted by the frequent formation of hybrid populations; this could suggest that the reported cases of homoploid speciation, may simply reflect a geographical distribution conducive to the formation and isolation of hybrid population. The Italian Sparrow seems to fit this scenario remarkably well (Runemark et al., 2018).

The search for signs of hybrid speciation in very large populations, for example yeast, would be an exciting avenue for hybrid speciation research in the future. That is be because, firstly, multiple DMIs, both inter- and intraspecific, have been identified in yeast (Kao et al., 2010; Matute et al., 2010; Hou and Schacherer, 2016). In addition, yeast can be easily maintained in the lab and lends itself to powerful experimental approaches. the large population size and the possibility to create and maintain multiple replicates are two strong advantages of experimental-evolution approaches to potentially quantify the reciprocal sorting of DMIs.

One remarkable genetic incompatibility in yeast that could serve as a contributor to hybrid speciation was identified by Bui et al. (2015). Intriguingly, the phenotype of the incompatibility is an increased mutation rate, which could provide the fuel for adaptation to a new environment upon hybridization (Taddei et al., 1997). Intrinsic incompatibilities are usually deleterious, however in this specific case they could give a temporary advantage to the hybrid population. Following the formation of a hybrid population, the incompatible haplotype could spread in the population while the hybrid population is adapting to new (harsh) conditions. Such a mechanism could therefore be powerful in a colonization scenario. In the initial stage, the mutator phenotype might increase the possibility of recruiting adaptive mutations. Sorting of the incompatibility happens only at a later stage, which results in a synergy of forming genetic isolating barriers from the parental source populations while delaying the incompatibility-sorting phase until the population has adapted to its environment.

## Conclusion

The probability of hybrid speciation is subject to continuing debate (Schumer et al., 2015; Nieto Feliner et al., 2017; Schumer et al., 2018). The reciprocal sorting of parental incompatibilities has been proposed as one credible mechanism to achieve hybrid speciation. Our work legitimates the existing disagreement by demonstrating that the hybrid speciation probability via reciprocal parental incompatibility sorting is highly variable and dependent on the linkage architecture and the dominance type of the involved incompatibilities. Specifically, the linkage architecture determines not only whether hybrid speciation is achievable or not, but also whether equal or unequal initial proportions of the parental populations are favorable for hybrid speciation. In addition, we show that across all studied scenarios, intermediate recombination rates maximize the likelihood of reciprocal sorting; i.e., interactions on the same chromosome are favorable for hybrid speciation. Altogether, we arrive at the prediction that in nature, hybrid speciation via reciprocal sorting of incompatibilities should indeed be rare; at the same time however, it can become almost deterministic (and, thus, repeatable) under specific genetic and demographic circumstances. Such circumstances could be met in microorganisms such as yeast, which have large population sizes and potential for repeated hybridization.

## Methods

### Simulations

Simulations are implemented in C++ (available at https://gitlab.com/evoldyn/four-loci) and each run ended when the population was monomorphic.

## Acknowledgments

We thank Roger Butlin, Florian Clemente, Aaron Comeault, Ilse Höllinger, Molly Schumer and the members of the Bank lab for helpful discussion and comments on the manuscript.

## References

R. Abbott, D. Albach, S. Ansell, J. Arntzen, S. Baird, N. Bierne, J. Boughman, A. Brelsford, C. Buerkle, R. Buggs, et al. Hybridization and speciation. Journal of Evolutionary Biology, 26(2):229–246, 2013.

M. Arnold, M. Bulger, J. Burke, A. Hempel, and J. Williams. Natural hybridization: how low can you go and still be important? Ecology, 80(2):371–381, 1999.

C. Bank, R. Bürger, and J. Hermisson. The Limits to Parapatric Speciation: Dobzhansky–Muller Incompatibilities in a Continent–Island Model. Genetics, 191(3):845–863, 2012.

N. Barton. The role of hybridization in evolution. Molecular Ecology, 10(3):551–568, 2001.

N. Barton and B. Bengtsson. The barrier to genetic exchange between hybridising populations. Heredity, 57(3):357–376, 1986.

W. Bateson. Heredity and variation in modern lights. Darwin and modern science, pages 85–101, 1909.

C. Buerkle, R. Morris, M. Asmussen, and L. Rieseberg. The likelihood of homoploid hybrid speciation. Heredity, 84(4):441–451, 2000.

D. Bui, E. Dine, J. Anderson, C. Aquadro, and E. Alani. A genetic incompatibility accelerates adaptation in yeast. PLoS genetics, 11(7):e1005407, 2015.

M. Colomé-Tatché and F. Johannes. Signatures of Dobzhansky–Muller Incompatibilities in the Genomes of Recombinant Inbred Lines. Genetics, 202(2):825–841, 2016.

A. Comeault. The genomic and ecological context of hybridization affects the probability that symmetrical incompatibilities drive hybrid speciation. Ecology and evolution, 8(5):2926–2937, 2018.

R. Corbett-Detig, J. Zhou, A. Clark, D. Hartl, and J. Ayroles. Genetic incompatibilities are widespread within species. Nature, 504(7478):135–137, 2013.

J. Coyne and H. Orr. Speciation. Sinauer Associates Sunderland, MA, 2004.

T. Dobzhansky. Studies on hybrid sterility. II. Localization of sterility factors in *Drosophila pseudoobscura* hybrids. Genetics, 21(2):113, 1936.

J. Feder, S. Egan, and P. Nosil. The genomics of speciation-with-gene-flow. Trends in Genetics, 28(7):342–350, 2012.

B. Gross. Genetic and phenotypic divergence of homoploid hybrid species from parental species. Heredity, 108(3):157, 2012.

R. Guerrero, C. Muir, S. Josway, and L. Moyle. Pervasive antagonistic interactions among hybrid incompatibility loci. PLoS Genetics, 13(6):e1006817, 2017.

J. B. S. Haldane. A mathematical theory of natural and artificial selection, part v: selection and mutation. In Mathematical Proceedings of the Cambridge Philosophical Society, volume 23, pages 838–844. Cambridge Univ Press, 1927.

P. Hedrick. Adaptive introgression in animals: Examples and comparison to new mutation and standing variation as sources of adaptive variation. Molecular Ecology, 22(18):4606–4618, 2013.

J. Hermansen, S. Sæther, T. Elgvin, T. Borge, E. Hjelle, and G. Sætre. Hybrid speciation in sparrows I: Phenotypic intermediacy, genetic admixture and barriers to gene flow. Molecular Ecology, 20(18):3812–3822, 2011.

J. Hou and J. Schacherer. Negative epistasis: a route to intraspecific reproductive isolation in yeast? Current genetics, 62(1):25–29, 2016.

J. Kang, M. Schartl, R. Walter, and A. Meyer. Comprehensive phylogenetic analysis of all species of swordtails and platies (Pisces: *Genus Xiphophorus*) uncovers a hybrid origin of a swordtail fish, *Xiphophorus monticolus*, and demonstrates that the sexually selected sword originated in the ancestral lineage of the genus, but was lost again secondarily. BMC Evolutionary Biology, 13(1), 2013.

K. C. Kao, K. Schwartz, and G. Sherlock. A genome-wide analysis reveals no nuclear Dobzhansky-Muller pairs of determinants of speciation between S. cerevisiae and S. paradoxus, but suggests more complex incompatibilities. PLoS genetics, 6(7):e1001038, 2010.

M. Kimura. On the probability of fixation of mutant genes in a population. Genetics, 47(6):713, 1962.

S. Lamichhaney, F. Han, M. Webster, L. Andersson, B. Grant, and P. Grant. Rapid hybrid speciation in Darwin’s finches. Science, 359(6372):224–228, jan 2018.

P. Larsen, M. Marchan-Rivadeneira, and R. Baker. Natural hybridization generates mammalian lineage with species characteristics. Proceedings of the National Academy of Sciences, 107(25):11447–11452, 2010.

J. Mallet. Hybrid speciation. Nature, 446(7133):279–283, 2007.

S. Marsit, J. Leducq, E. Durand, A. Marchant, M. Filteau, et al. Evolutionary biology through the lens of budding yeast comparative genomics. Nature Reviews Genetics, 18(10):581–598, 2017.

D. Matute, I. Butler, D. Turissini, and J. Coyne. A Test of the Snowball Theory for the Rate of Evolution of Hybrid Incompatibilities. Source: Science, New Series, 329(5998):1518–1521, 2010.

Je. Mavárez, C. Salazar, E. Bermingham, C. Salcedo, C. Jiggins, and M. Linares. Speciation by hybridization in heliconius butterflies. Nature, 441(7095):868–871, 2006.

H. Muller. Isolating mechanisms, evolution and temperature. In Biol. Symp, volume 6, pages 71–125, 1942.

G. Nieto Feliner, I. Álvarez, J. Fuertes-Aguilar, M. Heuertz, I. Marques, F. Moharrek, R. Piñeiro, R. Riina, J. Rosselló, P. Soltis, and I. Villa-Machío. Is homoploid hybrid speciation that rare? An empiricist’s view. Heredity, 118(6):513–516, 2017.

J. Ono, A. Gerstein, and S. Otto. Widespread Genetic Incompatibilities between First-Step Mutations during Parallel Adaptation of *Saccharomyces cerevisiae* to a Common Environment. PLoS biology, 15(1):e1002591, 2017.

H. Orr. A mathematical model of Haldane’s rule. Evolution, 47(5):1606–1611, 1993.

H. Orr. The population genetics of speciation: the evolution of hybrid incompatibilities. Genetics, 139(4):1805–1813, 1995.

D. Presgraves. A fine-scale genetic analysis of hybrid incompatibilities in *Drosophila*. Genetics, 163(3):955–972, 2003.

L. Rieseberg. Hybrid Origins of Plant Species. Annual Review of Ecology and Systematics, 28(2):359–389, 1997.

A. Runemark, C. Trier, F. Eroukhmanoff, J. Hermansen, M. Matschiner, M. Ravinet, T. Elgvin, and G. Sætre. Variation and constraints in hybrid genome formation. Nature Ecology & Evolution, 2018.

S. Scarpino, P. Hunt, F. Garcia-De-leon, T. Juenger, M. Schartl, and M. Kirkpatrick. Evolution of a genetic incompatibility in the genus xiphophorus. Molecular Biology and Evolution, 30(10):2302–2310, 2013.

M. Schumer, G. Rosenthal, and P. Andolfatto. How common is homoploid hybrid speciation? Evolution, 68(6):1553–1560, 2014.

M. Schumer, R. Cui, G. Rosenthal, and P. Andolfatto. Reproductive isolation of hybrid populations driven by genetic incompatibilities. PLoS Genet, 11(3):e1005041, 2015.

M. Schumer, G. Rosenthal, and P. Andolfatto. What do we mean when we talk about hybrid speciation? Heredity, pages 11–31, 2018.

D. Schwarz, B. Matta, N. Shakir-Botteri, and B. McPheron. Host shift to an invasive plant triggers rapid animal hybrid speciation. Nature, 436(7050):546–549, 2005.

O. Seehausen. Conditions when hybridization might predispose populations for adaptive radiation. Journal of Evolutionary Biology, 26(2):279–281, 2013.

O. Seehausen, G. Takimoto, D. Roy, and J. Jokela. Speciation reversal and biodiversity dynamics with hybridization in changing environments. Molecular Ecology, 17(1):30–44, 2008.

M. Servedio, J. Hermisson, and G. Doorn. Hybridization may rarely promote speciation. Journal of evolutionary biology, 26(2):282–285, 2013.

F. Taddei, M. Radman, J. Maynard-Smith, B. Toupance, PH. Gouyon, and B. Godelle. Role of mutator alleles in adaptive evolution. Nature, 387(6634):700, 1997.

M. Turelli and H. Orr. Dominance, epistasis and the genetics of postzygotic isolation. Genetics, 154(4):1663–79, 2000.

D. Turissini, J. McGirr, S. Patel, J. David, and D. Matute. The rate of evolution of postmating-prezygotic reproductive isolation in Drosophila. Molecular Biology and Evolution, 35(2):312–334, 2017.

S. Via. Divergence hitchhiking and the spread of genomic isolation during ecological speciation-with-gene-flow. Philosophical Transactions of the Royal Society B: Biological Sciences, 367(1587):451–460, 2012.

C.I. Wu. The genic view of the process of speciation. Journal of Evolutionary Biology, 14(6):851–865, 2001.

S. Yakimowski and L. Rieseberg. The role of homoploid hybridization in evolution: A century of studies synthesizing genetics and ecology. American Journal of Botany, 101(8):1247–1258, 2014.

S. Yeaman. Genomic rearrangements and the evolution of clusters of locally adaptive loci. Proceedings of the National Academy of Sciences, 110(19):E1743–E1751, 2013.

